# Flnc: Machine Learning Improves the Identification of Novel Full-length Long Noncoding RNAs from RNA Sequencing Data Without Transcriptional Initiation Profiles

**DOI:** 10.1101/2022.08.02.502545

**Authors:** Zixiu Li, Peng Zhou, Euijin Kwon, Katherine Fitzgerald, Zhiping Weng, Chan Zhou

## Abstract

Long noncoding RNAs (lncRNAs) play critical regulatory roles in human development and disease. However, many lncRNAs have yet to be annotated. The conventional approach to identifying novel lncRNAs from RNA sequencing (RNA-seq) data is to find transcripts without coding potential. This approach has a false discovery rate of 30-75%. The majority of these misidentified lncRNAs are RNA fragments or transcriptional noise and lack defined transcription start sites, which are marked by H3K4me3 histone modifications. Therefore, the accuracy of lncRNA identification can be improved by incorporating H3K4me3 chromatin immunoprecipitation sequencing (ChIP-seq) data. However, because of cost, time, and limited sample availability, most RNA-seq data lacks such data. This paucity of H3K4me3 data greatly hinders the efforts to accurately identify novel lncRNAs. To address this problem, we have developed software, Flnc, to identify both novel and annotated full-length lncRNAs from RNA-seq data without H3K4me3 profiles. Flnc integrates machine learning models built incorporating four types of features: transcript length, promoter signature, multiple exons, and genomic location. Flnc achieves state-of-the-art prediction power with an AUROC score over 0.92. Flnc significantly improves the prediction accuracy from less than 50% using the conventional approach to over 85%. Flnc is available via *https://github.com/CZhouLab/Flnc*.

## INTRODUCTION

Only about 1% of the human genome and less than 3% of RNA transcripts encode proteins(1). Long noncoding RNAs (lncRNAs) are a subset of noncoding RNAs longer than 200 nucleotides. Similar as messenger RNAs (mRNAs), lncRNAs contain 5′ caps and 3′ polyadenylation (polyA) tails. LncRNAs play regulatory roles in numerous biological processes, including human stem cell development and immunity (2–9), and they have been implicated in many diseases, including neurological disorders, cardiovascular, lung, and liver diseases, infectious diseases, and cancer (8, 10, 11). LncRNAs are also associated with complex genetic traits, playing important roles in gene regulation by functioning as protein scaffolds, guiding DNA-protein interactions, controlling post-transcriptional regulation, and functioning as cis-regulatory elements at enhancers (12). However, because evolutionary constraints in noncoding regions are lower than in coding regions, lncRNAs are less conserved and evolve faster than coding genes (13). Increasingly, studies have shown that lncRNAs exhibit striking disease-specific expression patterns and are potential drivers and modifiers of disease (9, 14). Current databases like GENCODE (15), NONCODE (16), and LNCipedia (17) lack annotations for many disease-or cell-type-specific lncRNAs. For example, we previously found that about 40% of lncRNAs expressed in human hepatic stellate cells are not yet included in the database (18). Therefore, it is critical to create a universal tool for the identification of novel lncRNAs which have not been annotated in the current databases, and this tool can be applied across various diseases and developmental processes.

Advances in high-throughput sequencing techniques have produced a large amount of publicly available RNA sequencing data from various tissues, cell lines, and disease models. To date, there are more than 12,700 study series in the NCBI Gene Expression Omics (GEO) database with available RNA-seq datasets. This large amount of data offers great opportunities to identify novel lncRNAs expressed in specific biological samples. In addition, the rapid development of computational methods for analyzing sequencing data including methods for transcript assembly (e.g., Scripture (19), Trinity (20), Cufflink (21), StringTie (22), Strawberry (23), and TransComb (24)) and methods for examining the coding abilities of transcripts (e.g., CPAT (25), LGC (26), PLEK (27), and CPPred (28)), have allowed researchers to generate catalogs of putative lncRNAs—assembled transcripts without protein-coding potential. However, these methods cannot distinguish between true lncRNAs and false-positive lncRNAs, most of which are nonfunctional transcribed fragments and transcriptional noise. Unlike the false lncRNAs, true lncRNAs are high-confidence full-length lncRNA transcripts that include transcriptional start sites (TSSs). In addition, lncRNAs have low expression levels compared to mRNAs, so we cannot distinguish true lncRNA from false lncRNAs purely based on expression levels.

To identify true lncRNAs, including the novel lncRNAs, some studies have focused on identifying lncRNAs that are conserved across mammals (29, 30). These methods have improved accuracy but may fail to detect human-specific lncRNAs (31). To improve the identification of true lncRNAs, especially the novel lncRNAs, markers of transcription initiation have been incorporated into identification pipelines. The trimethylation of histone H3 at lysine 4 (H3K4me3) is a chromatin modification known to mark transcription start sites of active genes (32), including lncRNA genes.

Therefore, H3K4me3 profiling data (ChIP-seq or CUT-RUN seq data) are commonly used to identify lncRNAs. Unfortunately, over 96% of data series in the NCBI GEO database with available RNA-seq data lack matching H3K4me3 profile data. Therefore, identifying lncRNAs from these GEO data series and their associated samples is not reliable. Due to cost, time, limited sample availability generating H3K4me3 profiles is often impractical; this is especially true for patient-derived clinical samples. When transcription initiation profiling data is sparse, up to 75% of lncRNAs identified by existing methods will be transcribed fragments or background transcription noise, not true lncRNAs (**Suppl Fig 1B**).

Therefore, we sought to improve the accuracy of novel lncRNA identification from RNA-seq data lacking matched H3K4me3 ChIP-seq. We developed a machine learning-based method to distinguish between true and false lncRNAs. Incorporating this new method, we developed *Flnc*, software that can directly identify true lncRNAs, including novel and annotated lncRNAs, from RNA-seq data. We assessed *Flnc* using over 240,000 putative lncRNAs from 46 published RNA-seq datasets that had matching H3K4me3 ChIP-seq data. In five independent test datasets, the *Flnc* pipeline significantly reduced the rate of false positives and achieves an over 85% prediction accuracy which significantly outperform the conventional method with 55% of prediction accuracy. The *Flnc* pipeline integrates pre-processing, mapping, transcript assembly, evaluation of protein-coding ability, and evaluation of transcripts using our machine-learning algorithm. The *Flnc* pipeline, which is implemented on the user-friendly Singularity platform, has minimal pre-requisites and is easily portable. The *Flnc* can also run with parallelize tasks to optimizes computing resources.

## MATERIAL AND METHODS

### Collection of sequencing datasets to generate benchmark lncRNAs

Generating a benchmark dataset of true and false lncRNAs requires stranded polyA-selected RNA-seq and sample-matched H3K4me3 ChIP-seq. We identified 388 data series with the keywords “RNA seq,” “H3K4me3,” and “Homo sapiens” from the NCBI GEO database through February of 2021. From among the 388 GEO studies, we selected 61 datasets with available sample-matched, stranded polyA-selected RNA-seq and H3K4me3 ChIP-seq data.

For the 61 datasets, we examined the quality of RNA-seq and ChIP-seq data and removed 15 datasets, which had poor RNA-seq or ChIP-seq data quality. We considered the quality of RNA-seq data poor if more than half of the reads had QC scores less than 35. For ChIP-seq data, we considered their quality poor if the number of called peaks in the dataset was an outlier among peak numbers found in the 61 datasets (extremely small <2000 or extremely large >200,000). This process resulted in 46 datasets. The RNA-seq and ChIP-seq data within each of the selected 46 high-quality datasets were generated from the same type of sample. The datasets are heterogenous, including both single-end (50%) and paired-end (50%) RNA-seq data and each dataset has one to four RNA-seq replicates. We indexed these datasets 1 to 46 in chronological order by submission date and used them to identify true and false lncRNAs in each dataset as benchmark data.

After identifying true and false lncRNAs in each dataset (see below), we split the benchmark dataset into training and testing sets based on the submission dates of the RNA-seq data into the NCBI GEO database (33). The training set included the 41 datasets that were submitted to the GEO database before 2019; we used the 5 datasets generated in 2019 and 2020 as the testing set.

### Identification of putative lncRNAs

First, we mapped the RNA-seq data to the human reference genome (hg38) using HISAT2 v 2.0.5 (34) and assembled transcripts using both StringTie v1.3.4 (22) and Strawberry v1.1.2 (23). For each sample, we then merged assembled transcripts with the StringTie merge function. Next, we examined the coding potential of the assembled transcripts using CPAT v1.2.4 (25), LGC v1.0 (26), PLEK v1.2. (27), and CPPred (28). We excluded the transcripts with coding potential defined in any of the above tools. Next, we excluded the resulting transcripts as well as any transcripts that overlapped protein-coding genes or pseudogenes on the same strand. We also excluded transcripts overlapping other annotated noncoding RNAs, including snoRNA, rRNA, tRNA and microRNAs on the same strand. From the remaining transcripts, we selected expressed transcripts that were over 200 nucleotides long as putative lncRNAs. (See **Supplemental Methods** for details.)

### Identification of H3K4me3 peaks using H3K4me3 ChIP-seq data

We aligned H3K4me3 ChIP-seq reads to the human reference genome (hg38/GRCh38) using the BWA v0.7.5a toolkit (35). For reads with mean read length ≥ 70 bp, we used BWA-MEM, and for short reads, we used BWA-aln. Next, we used the MACS2 v2.2.7.1 (36) peakcall function to identify H3K4me3 as previously (18) with the following settings: -q 0.01 –broad –broad-cutoff=0.01 –nomodel –extsize 300. For H3K4me3 ChIP-seq data without matched control data, MACS2 called peaks based on the H3K4me3 input ChIP-seq data. For H3K4me3 ChIP-seq data with matched control data, MACS2 called peaks by comparing the bam files to the matched background control. We used the H3K4me3 broad peaks called by MACS2 as transcription initiation markers (37, 38).

### Identification of true and false lncRNAs based on H3K4me3 ChIP-seq data

We used our previously established approach (18) to identify true and false lncRNAs among putative lncRNAs. Because H3K4me3 chromatin modification is well known as transcriptional initiation marks of active genes (32), including lncRNA genes, we examined the distance between the 5′ ends of putative lncRNAs and the matched H3K4me3 peaks. If the H3K4me3 peak was within 1kb of a putative lncRNA, we considered it a true lncRNA; otherwise, we considered it false.

### Normalization of transcript lengths

We calculated the length of putative lncRNAs by summing the length of all the transcript′s exons. Next, we log-transformed the lengths. We calculated the upper and lower limits as mean plus 3x standard deviation and mean minus 3x standard deviation, respectively. Then, we set the outlier data points of the log-transformed lengths as the value of upper or lower limits. Finally, we scaled the log-transformed values of transcript lengths for each putative lncRNA into the range between 0 and 1 using the min-max normalization technique.

### Identification of promoter regions

We used TSSG software (39) to identify the potential promoters in the genomic regions ± 1kb of the 5′ end of putative lncRNAs. TSSG detects promoters by scanning for transcription factor binding sites and is considered to be one of the most accurate mammalian promoter prediction programs with the fewest false positive predictions (40).

### Classification of putative lncRNAs by genomic location

We used the genomic locations of protein-coding genes annotated by the GENCODE Project release 29 (41), and compared them to the genomic locations of putative lncRNAs using the BEDTools suite and GffCompare tool (43). We classified putative lncRNAs—true and false, into three categories by genomic locations (18, 44): divergent, antisense, and intergenic. A putative lncRNA was classified as divergent if the 5′ end of its locus was within ± 2 kb of the TSS of a protein-coding gene on the opposite strand (18, 44). Putative lncRNAs that were antisense to protein-coding genes and overlapped the gene by at least one base pair were classified as antisense. All remaining putative lncRNAs were classified as intergenic.

### Calculation of the exon numbers of putative lncRNAs

The number of exons of putative lncRNAs were counted based on the assembled transcripts in GTF format, which include the information of exon numbers and coordinates of each exon.

### Measurement feature importance

To determine the feature importance for all machine learning models, we used the permutation approach (45). We scaled the importance scores into the range between 0 and 1 by dividing each importance score by the importance score of the most significant feature.

### Collection of other RNA-seq datasets and matched H3K4me3 ChIP-seq data

To evaluate *Flnc*′s performance on other types of RNA-seq data, we collected four additional datasets, each corresponding to one sample from the GEO database. These four datasets all contained RNA-seq data and matched H3K4me3 ChIP-seq. Two datasets included stranded rRNA-depleted RNA-seq (accession numbers: GSE179184 and GSE189861), and two included unstranded polyA-selected RNA-seq data (accession numbers: GSE72131 and GSE58740).

## RESULTS

### Generation of a benchmark dataset of true and false lncRNAs

To benchmark our new tool, we first updated our previous computational pipeline (18, 44) (**Fig 1A**, see **Materials and Methods**). This pipeline distinguishes between true and false lncRNAs by first identifying potential lncRNAs from raw RNA-seq data, then examining if putative lncRNAs have H3K4me3 peaks near their 5′ ends. When we applied this pipeline to the 46 publicly available datasets with matched RNA-seq and ChIP-seq data, we identified the hundreds to the thousands true and false lncRNAs in each individual dataset (**Suppl Fig 1A**). Each putative lncRNA found within every dataset was counted as one data point, then it totally resulted in 95,265 true lncRNA data points and 149,147 false lncRNA data points. These true and false lncRNA data points constitute our benchmark dataset.

**Figure 1.**
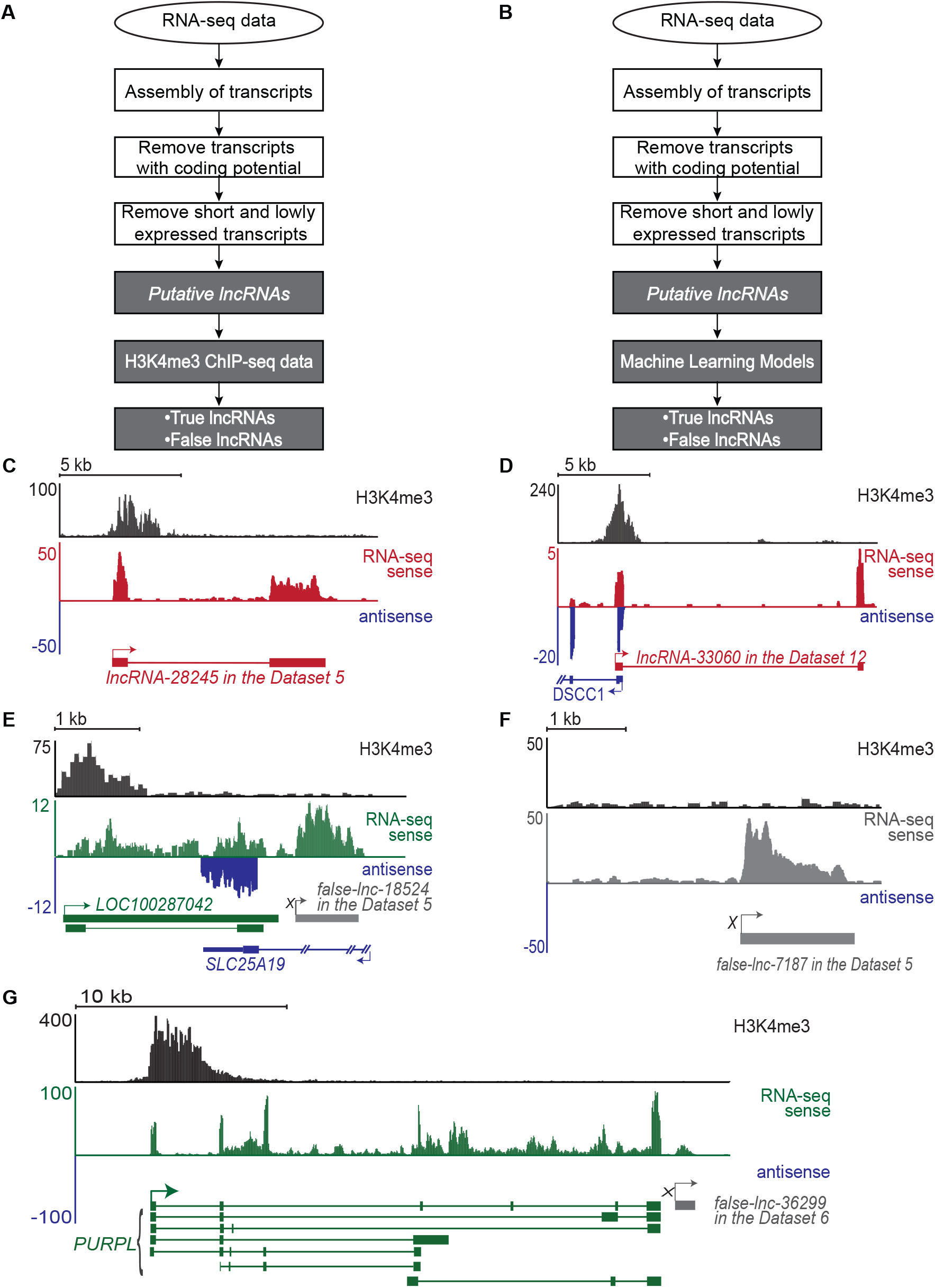
Identification of true and false lncRNAs. (A) Flowchart of the standard computational pipeline for identifying true and false lncRNAs (see Materials and Methods for details). The identified true lncRNAs include both novel and annotated lncRNAs. The standard pipeline requires both RNA-seq data and H3K4me3 ChIP-seq data generated from the same sample. (B) Flowchart of *Flnc* pipeline. *Flnc* integrates machine learning models to identify true and false lncRNAs directly from RNA-seq data, without matched H3K4me3 profiles. The *Flnc* pipeline includes two steps: first, *Flnc* identifies putative lncRNAs from RNA-seq data (see Materials and Methods for details). Then, *Flnc* classifies putative lncRNAs as true or false based on built-in machine learning models. The true lncRNAs predicted by *Flnc* include both novel and annotated lncRNAs. (C) A novel true lncRNA detected in Dataset 5. lncRNA-28245 (red) is located in an intergenic region on chromosome 3. The characteristic H3K4me3 peak identifies it as a true lncRNA. (D) A novel true lncRNA identified in Dataset 12. lncRNA-33060 (red) is located on the sense strand and transcribed divergently from the promoter of the protein-coding DSCC1 (blue) gene. (E) A false lncRNA identified in Dataset 5. False-lnc-18524.1 (grey) is located downstream of LOC100287042 (green) and antisense to SLC25A19 (blue). This false lncRNA is supported by an abundance of RNA-seq reads but lacks H3K4me3 peaks at the 5′ end. The false-lncRNA could be a fragment of an isoform transcript of the LOC100287042 gene. (F) An intergenic false lncRNA identified in Dataset 5. This lncRNA is supported by an abundance of RNA-seq reads but lacks a H3K4me3 peak at the 5′ end. (G) A false lncRNA identified in Dataset 6. False-lnc-36299 (grey) is downstream of the protein-coding gene PURPL (green). Compared to PURPLE, false-lnc-36299 is expressed at relatively low levels; therefore, it may be the result of RNA polymerase continuing beyond the polyA signal sequences when transcribing PURPLE (transcriptional noise) (Eaton and West, 2018; Luo and Bentley, 2004). For each lncRNA track, H3K4me3 peaks mark the site of transcription initiation (black, top). RNA-seq reads supporting true lncRNA transcripts are shown in red (RNA-seq sense). RNA-seq reads supporting annotated genes in the sense strand are shown in green (RNA-seq sense). Antisense transcripts are shown in blue. RNA-seq reads supporting false lncRNAs are shown in gray or in green when these reads continue the RNA-seq reads of supporting protein-coding transcripts. The genomic structure for each gene is shown below the RNA-seq tracks. True lncRNAs are shown in red; false lncRNA are shown in grey; annotated genes in the antisense strand are shown in blue, and annotated genes in the sense strand are shown in green. Boxes represent exons, lines represent introns, and arrows represent the start and direction of transcription. Two annotated genes, LOC100287042 and PURPL, have multiple isoforms. True lncRNAs identified in this study are named with the prefix “lncRNA” followed by the locus number assigned during assembly. False lncRNAs are named with the prefix “false-lnc” followed by the locus number assigned during assembly.

Using our pipeline, we identified 43,941 true lncRNAs unannotated in GENCODE, which are supported by RNA-seq and H3K4me3 ChIP-seq data. Examples of true lncRNA identified by the pipeline are illustrated in **Fig 1C-D**. However, for 61% of the putative lncRNAs from the benchmark dataset, we could not locate a transcription start site, as indicated by H3K4me3 signal. Using these criteria, we determined that 30%-75% of the putative lncRNAs in each dataset were not true lncRNAs (**Fig 1E-G and Suppl Fig 1B**). Therefore, without H3K4me3 ChIP-seq data, 30%-75% of putative lncRNAs identified from RNA-seq data alone can be expected to be false hits.

### Four genomic features can be used to distinguish true and false lncRNAs

To improve identification of lncRNAs from RNA-seq data lacking matched H3K4me3 ChIP-seq data, we developed a new computational tool, *Flnc* (**Fig 1B**). *Flnc* integrates four types of genomic features of lncRNAs into the machine learning models: transcript length, promoter signature, multiple exons, and genomic location. We hypothesized that these features would provide enough information to distinguish between true and false lncRNAs.

Most false lncRNAs are transcript fragments or transcriptional noise. Therefore, we hypothesized that true lncRNAs would tend to be longer than false lncRNAs. To test this hypothesis, we examined the transcript length of all putative lncRNAs in the benchmark dataset. Read length and sequencing depth differ across various RNA-seq data sets and these factors affect transcript assembly (46–48). To control for these differences, for each of the 46 datasets, we normalized the transcript length of putative lncRNAs to a value between 0 and 1 (see **Materials and Methods**). After normalization, the transcript lengths of true lncRNAs were significantly longer than those of false lncRNAs (**Fig 2A and Suppl Fig 2A**).

**Figure 2.**
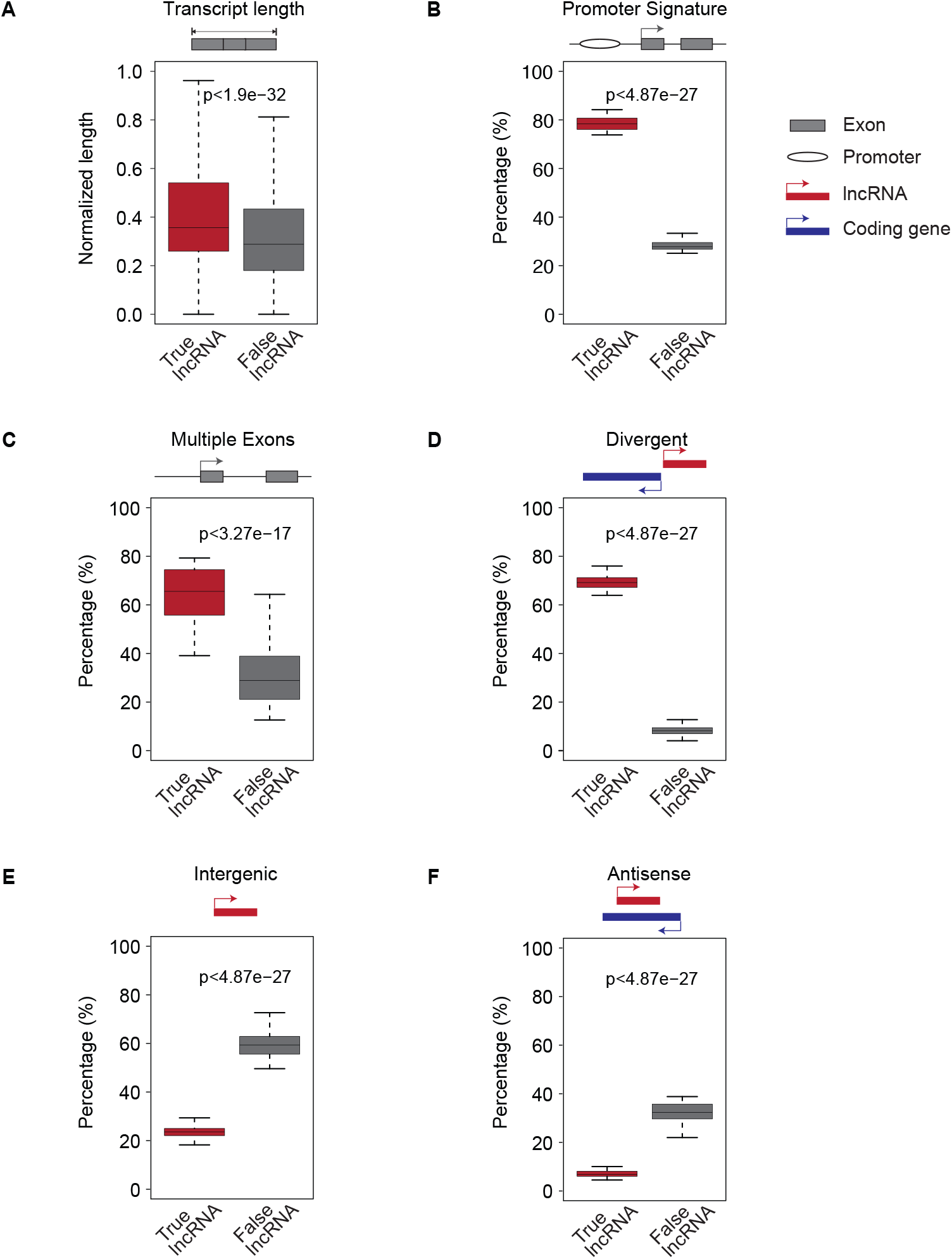
Four features exhibit significant differences between true and false lncRNAs in the 46 benchmark datasets. True lncRNAs can be distinguished from false lncRNAs because they tend to be longer (A), to have a predicted upstream promoter signature (B), and to be more likely to have multiple exons (C). They can also be distinguished by genomic location and are more likely to be divergently transcribed from the promoter of a protein-coding gene (D). True lncRNAs are less likely to be in intergenic regions (E) or antisense (F) to a protein-coding gene. For the graph (A), the boxplot represents the scaled transcript lengths of true and false lncRNAs. For the graphs (B-F), each boxplot represents the percentage of a feature of true and false lncRNAs among the identified putative lncRNAs for each of the 46 benchmark datasets. P values were calculated by two-sided Wilcoxon-Mann-Whitney test.

Whereas RNA fragments and transcripts resulting from noisy transcription lack promoters and TSSs, true lncRNAs should have upstream promoters. Therefore, we used TSSG software (39) to identify putative promoters upstream of true and false lncRNAs and examined the percentage of true and false lncRNAs with upstream promoter signatures. Almost 80% of true lncRNAs had putative upstream promoters. In contrast, we find this feature in only about 20% of false lncRNAs (**Fig 2B and Suppl Fig 2B**).

Because the vast majority of single-exon transcripts result from transcriptional and alignment noise (49, 50), we hypothesized that false lncRNAs (transfrags or noise) would have fewer exons than true lncRNAs. To test this hypothesis, we examined the percentage of multiple exon transcripts among true and false lncRNAs. We found that true lncRNAs are significantly more likely to have multiple exons than false lncRNAs (p-value < 3.27e-17; **Fig 2C and Suppl Fig 2C**).

Over 60% of lncRNAs have been shown to be divergently transcribed from the promoter regions of protein-coding genes (18, 44). This suggests that genomic context could be used to distinguish between true and false lncRNAs. We examined the true and false lncRNAs in our benchmark dataset, classifying them into three categories—divergent, antisense, and intergenic, based on their genomic locations. Consistent with previous findings, more than 60% of true lncRNAs were divergent transcripts of protein-coding genes, while less than 20% of false lncRNAs in each of the 46 datasets were divergent transcripts (**Fig 2D and Suppl Fig 2D**). Additionally, a small fraction of true lncRNAs originate from intergenic regions or are antisense to coding genes. In contrast, about 60% of false lncRNAs are in intergenic regions and 30% are antisense to coding genes (**Fig 2E-F and Suppl Fig 2E-F**). Therefore, true and false lncRNAs show significantly different patterns in terms of genomic location.

In conclusion, true and false lncRNAs show significant differences in these four features and these features can be used to distinguish between the two. Therefore, we proceeded to integrate these features into our machine learning models.

### Multiple machine learning models can distinguish between true and false lncRNAs

We trained seven of the most common machine learning (ML) algorithms using 81,420 true and 123,764 false lncRNA data points from the training set. These putative lncRNAs were identified from 41 datasets submitted to the GEO database before 2019. The seven ML algorithms include: logistic regression (LR), k-nearest neighbors (KNN), decision tree (DT), random forest (RF), naïve Bayes (NB), linear kernel support vector machine (SVM), and radial-basis-function (RBF) kernel SVM.

To fit and select the best model for each ML algorithm, we first used the 10-fold cross validation approach to train and evaluate each model using all possible combinations of hyperparameters (**Fig 3A**). We divided the putative lncRNAs in the training set randomly into ten non-overlapping subsets (folds). For each ML algorithm, we held one of the ten subsets of putative lncRNAs aside and trained a model using each set of hyperparameter values on the other nine subsets. We then evaluated the performance of the model on the remaining held-aside data. We repeated the process ten times, each time holding aside a different subset of data. We considered the hyperparameter values that defined the model with the best mean F1 score—the harmonic mean value of the precision and sensitivity—the optimal model architecture. Next, we trained the optimal model architecture on the entire training set to build the models for *Flnc*. On the training set, the random forest, decision tree and KNN models have the best overall prediction performance based on the F1 score and AUROC score (**Fig 3B**). The random forest model resulted in F1=0.84 and AUROC=0.94 values, the decision tree model resulted in F1=0.83 and AUROC=0.93 values, and the KNN model resulted in F1=0.82 and AUROC=0.92 values. Compared to the other models, these three models also have better accuracy, precision, and specificity. In contrast, the linear SVM is the worst performing of all models with respect to the F1 score, sensitivity, and accuracy.

**Figure 3.**
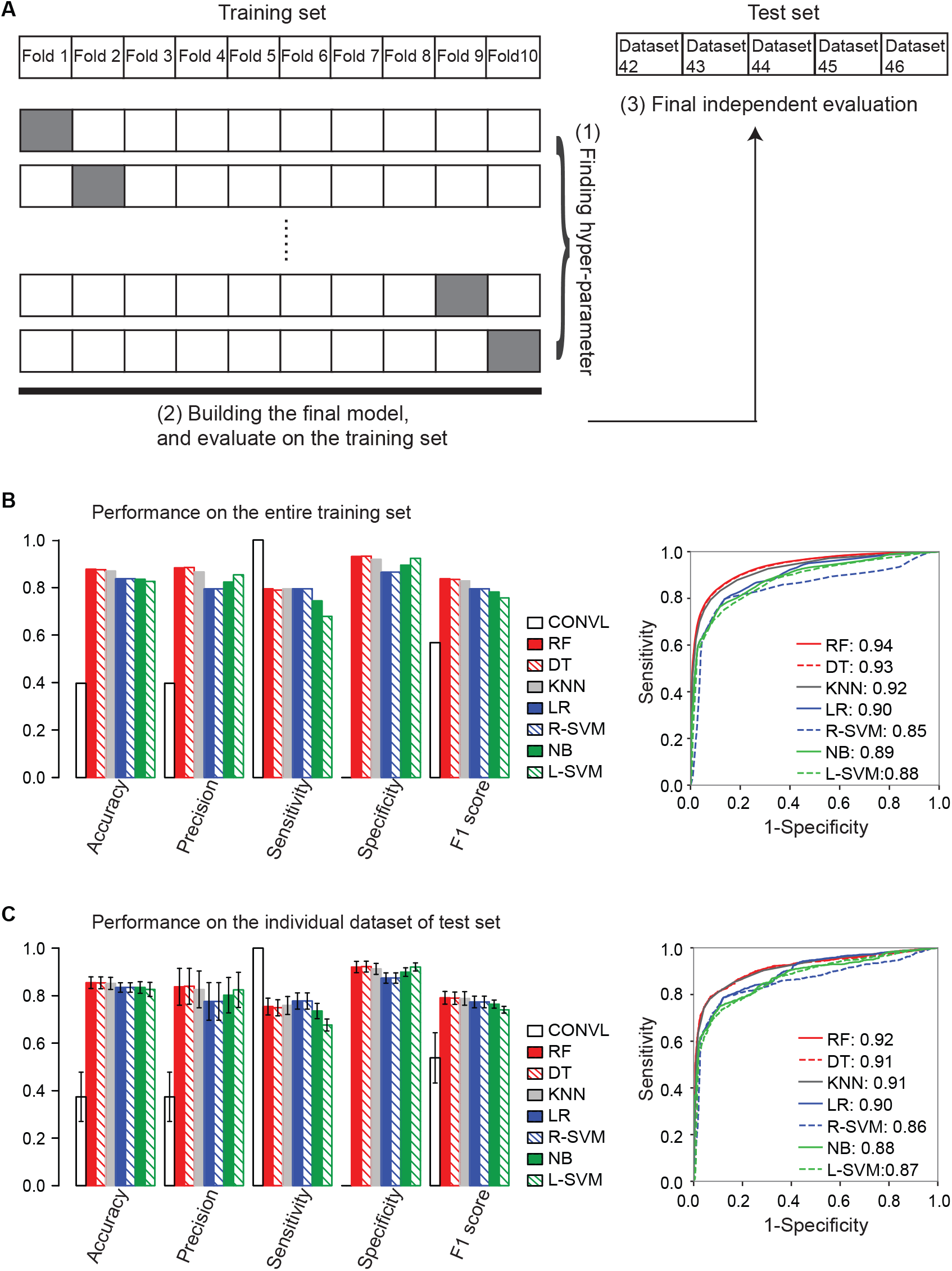
Machine learning model construction and evaluation. (A) The overall architecture of *Flnc* training and testing. The 81,420 true and 123,764 false lncRNA data points from the benchmark dataset were included in the training set. Each putative lncRNA found within every dataset were counted as one data point. These true and false lncRNAs were identified from the 41 datasets which were submitted to the GEO database before 2019. The putative lncRNAs of an additional five datasets that were generated in 2019 and 2020 were used as the test set. The best hyperparameters, specifically the hyperparameters yielding the best mean F1 score, were chosen using the ten-fold cross-validation approach as follows. The training set was randomly split into ten even folds, nine of which were used in training the models. The tenth fold was used for validation. We repeated this process holding each individual fold out and training with the other nine folds. From this, we derived ten models for each hyperparameter setting. For each of the ten models, we calculated the mean F1 score for each hyperparameter setting. After selecting the best hyperparameter setting, the final model was built using the entire training set and we examined the performance of the final model on the entire training set. Finally, we evaluated the performance of the final model on the test set, composed of the five independent datasets generated in 2019 and 2020 (Datasets 42-46). (B) The seven final models of *Flnc* outperformed the conventional method (without models) on the entire training set, and (C) on the five individual datasets of test set. The left bar graphs of (B-C) show the performance metrics accuracy, precision, sensitivity, specificity and F1 score. The right graphs (B-C) show the receiver operating characteristic (ROC) curve for each model. Area Under the ROC curve (AUROC) score is shown next to each ML model. The ROC curve of (C) is the ROC curve of the seven models on the dataset 46. Please see Supplemental Figure 3B for the ROC curves of the four additional test datasets (dataset 42-25). The seven models are ranked by the F1 score from largest to smallest. *Abbreviations: CONVL, conventional approach; RF, random forest; DT, decision tree; R-SVM, RBF support vector machine; L-SVM, linear support vector machine; LR: linear regression; NB, naïve bayes; KNN*, k-nearest neighbors.

To demonstrate *Flnc*′s ability to predict true lncRNAs from independent RNA-seq data, we tested *Flnc* on a test set composed of five independent datasets that were released to the GEO database after 2019 (**Fig 3C**). These five datasets were generated from multiple biological samples, including the MOLM-13 human myeloid leukemia cell line, the HUDEP-2 erythroid cell line, the Jurkat leukemia cells, and the H1299 non-small cell lung cancer cell line. We evaluated the performance of *Flnc* both on the entire test set (**Suppl Fig 3A**) and on the five individual datasets within the test set (**Fig 3C and Suppl Fig 3B**). Consistent with the training set results, the random forest, decision tree and KNN models have the best overall prediction abilities as indicated by the F1 and AUROC scores (**Fig 3C and Suppl Fig 3B**), although the seven models achieve 72%-87% consistency in lncRNA prediction (**Suppl Fig 3C**). These three models achieve 93%-96% consistency in lncRNA prediction (**Suppl Fig 3D**) with the accuracy of 0.85 or greater and precision of 0.83 or more. As with the training set, based on the F1 score and accuracy, the linear SVM and naïve bayes models have the worst performance. Variations in performance between datasets are small (**Fig 3C**) with a standard deviation in F1 score of less than 0.03 and a standard deviation in AUROC score of less than 0.02.

Furthermore, the ensemble approach can further improve prediction precision from 83% to 87% and specificity from 92% to 95% at the cost of reducing the sensitivity (**Suppl Fig 3E**). The ensemble approach used here is that lncRNA will be taken as true if all seven models predict it as a true lncRNA.

### Many lncRNAs identified by *Flnc* are novel and are supported by H3K4me3 profiles

Because true lncRNAs include both novel and annotated lncRNAs, we examined the novel lncRNAs among the true lncRNAs predicted by *Flnc* in the five independent test datasets. We found that up to 60% of true lncRNAs predicted by *Flnc* have not yet been annotated in the GENCODE database (**Fig 4A**). Among these novel lncRNAs identified by *Flnc*, most can be verified by H3K4me3 ChIP-seq data (**Fig 4B**). Compared to the conventional method, a novel lncRNA identified by Flnc has almost double the chance of being confirmed by H3K4me3 (**Fig 4B**). Taking the set of novel predicted lncRNAs common to all seven models improved the chance of being confirmed even further (**Fig 4B**).

**Figure 4.**
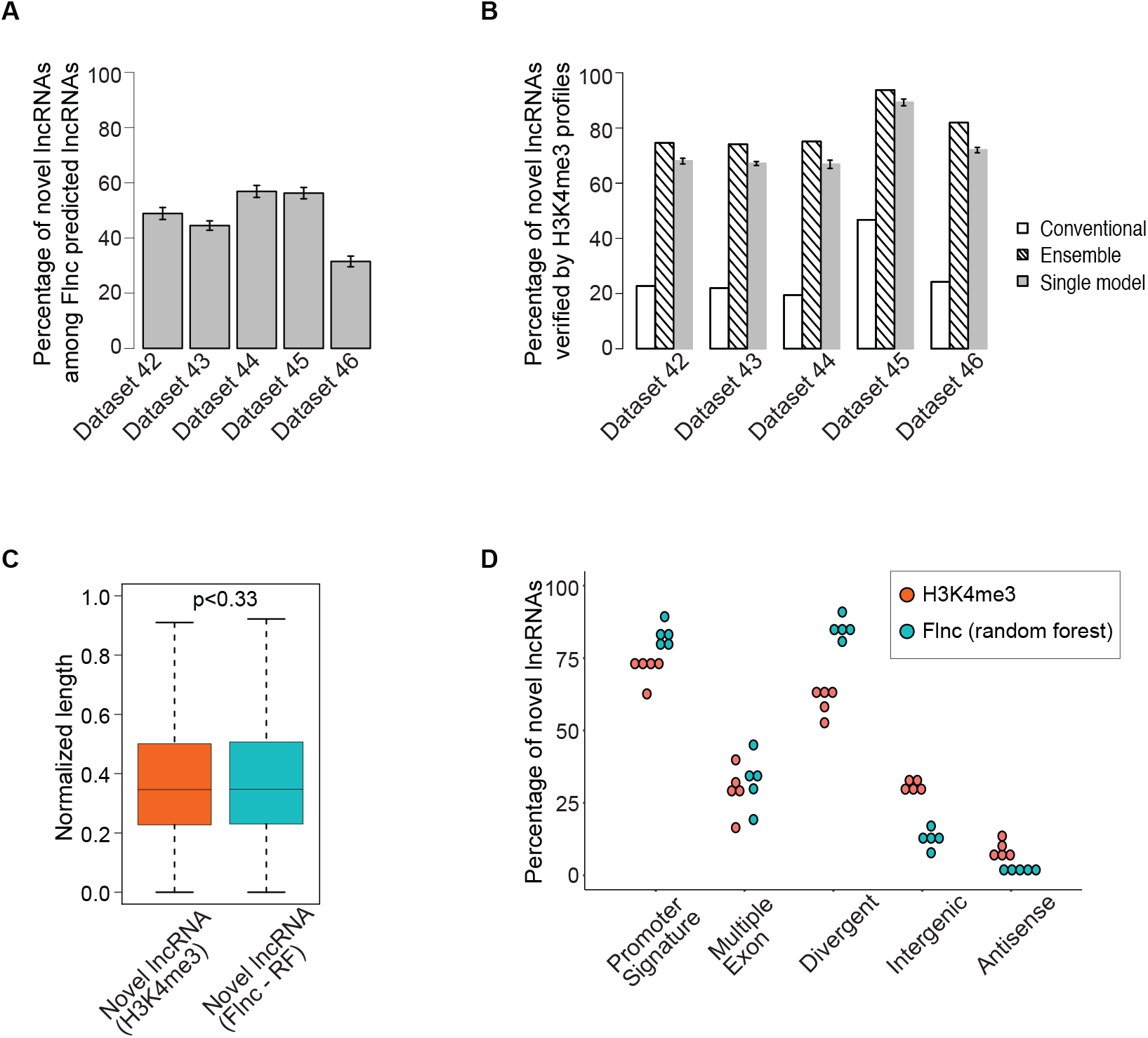
Most novel lncRNAs predicted by *Flnc* are supported by H3K4me3 profiles. (A) A large proportion of true lncRNAs predicted by *Flnc* are novel lncRNAs in the five individual test datasets. (B) The percentage of novel lncRNAs predicted by the conventional method without models and *Flnc* methods are verified by H3K3me3 profiles. Most novel lncRNAs predicted by the conventional method cannot be verified by H3K4me3 profiles, whereas most of the novel lncRNAs predicted by *Flnc* methods (*ensemble approach* or each of the seven single models) can be verified by H3K4me3 profiles. The error bar of the gray bar represents the standard deviation for the results of the seven ML models. (C)The novel lncRNAs predicted by *Flnc* (with random forest model) shows the similar normalized transcript length distribution as the novel lncRNAs determined by H3K4me3 ChIP-seq data. (D) The novel lncRNAs predicted by *Flnc* (with random forest model) includes significantly more divergent transcripts, and transcripts with promoter signatures than the novel lncRNAs determined by H3K4me3 ChIP-seq data, whereas multiple exon features exhibit similar percentage between these two groups of novel lncRNAs.

Next, we examined the genomic features of novel lncRNAs predicted by *Flnc* in the five independent test datasets. Because random forest has the best F1 score and AUROC score, we compared the genomic features of novel lncRNAs predicted with the random forest model with these of novel lncRNAs identified by the H3K4me3 ChIP-seq data (**Fig 4C-D**). The novel lncRNAs predicted by *Flnc* and those identified by H3K4me3 profiles were similar in terms of transcript length (**Fig 4C**) and multiple exons (**Fig 4D**), but the novel lncRNAs predicted by *Flnc* were more likely to have promoter signatures and to be divergent transcripts (**Fig 4D**). We observed similar trends among all the true lncRNAs predicted by Flnc and those determined by H3K4me3 profiles (**Suppl Fig 4A-B**).

### *Flnc* predicts true lncRNAs in multiple types of RNA-seq samples

For both the training and test data, we used stranded RNA-seq data generated from polyA-selected RNA. However, many RNA-seq datasets are generated not from poly-A RNA but from ribosomal-RNA (rRNA) depleted RNA, and some are sequenced without strand information. To evaluate the performance of *Flnc* in other types of RNA-seq data, we tested *Flnc* on two stranded RNA-seq datasets that generated using rRNA depletion (**Fig 5A**) and on two unstranded polyA RNA-seq datasets (**Fig 5B**).

**Figure 5.**
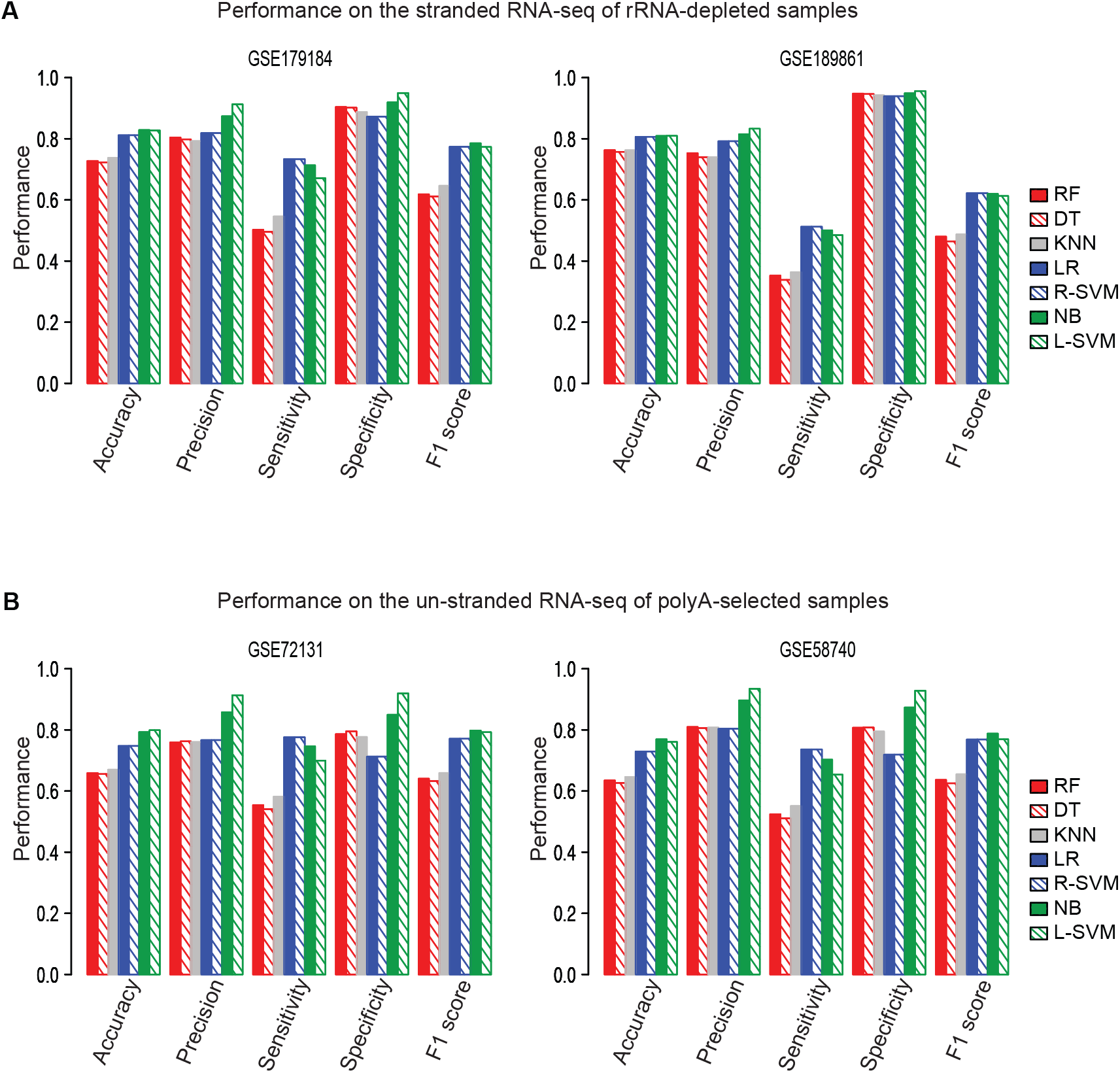
*Flnc* accurately identifies true lncRNAs from rRNA-depleted and unstranded RNA-seq datasets. (A) Performance of *Flnc* on two stranded RNA-seq datasets generated from ribosomal RNA (rRNA)-depleted samples (GSE179184 on the left, GSE189861 on the right). (B) Performance of *Flnc* on two un-stranded polyA-selected RNA-seq datasets (GSE72131 on the left, GSE58740 on the right).

On both stranded rRNA-depleted RNA-seq and unstranded polyA-selected RNA-seq data, four models—naïve Bayes, linear SVM, RBF SVM and logistic regression—have similar F1 scores and. Based on the F1 scores, these four models performed better than other three models. (**Fig 5**). On these types of data, the linear SVM model achieved the best prediction precision and accuracy; the naïve Bayes model ranks as the second best by the precision metric. While the random forest, decision tree and KNN models ranked as among the best models when run on polyA-selected RNA-seq data, their performance on rRNA-depleted RNA-seq data ranked as the worst by the metrics of accuracy, precision, sensitivity, and F1 score (**Fig 5**).

### Divergent transcription is the most important feature for predicting true lncRNAs

We evaluated the significance of the selected genomic features for each machine learning model. The feature importance score indicates the relative importance of each genomic feature (transcript length, promoter signature, multiple exons, and genomic location) for *Flnc* in detecting true lncRNAs from RNA-seq data. Because the genomic location feature includes three categories (divergent, antisense, intergenic), we evaluated the feature importance for six categories: transcript length, promoter signature, multiple exons, divergent transcript, antisense transcript, and intergenic transcript. We found that divergent transcript feature is the most important category across all machine learning models (**Fig 6**). For three models—random forest, decision tree, and KNN—promoter signature ranks as the second most important feature. In contrast, the intergenic and antisense transcript features are the least important in all models (**Fig 6**).

**Figure 6.**
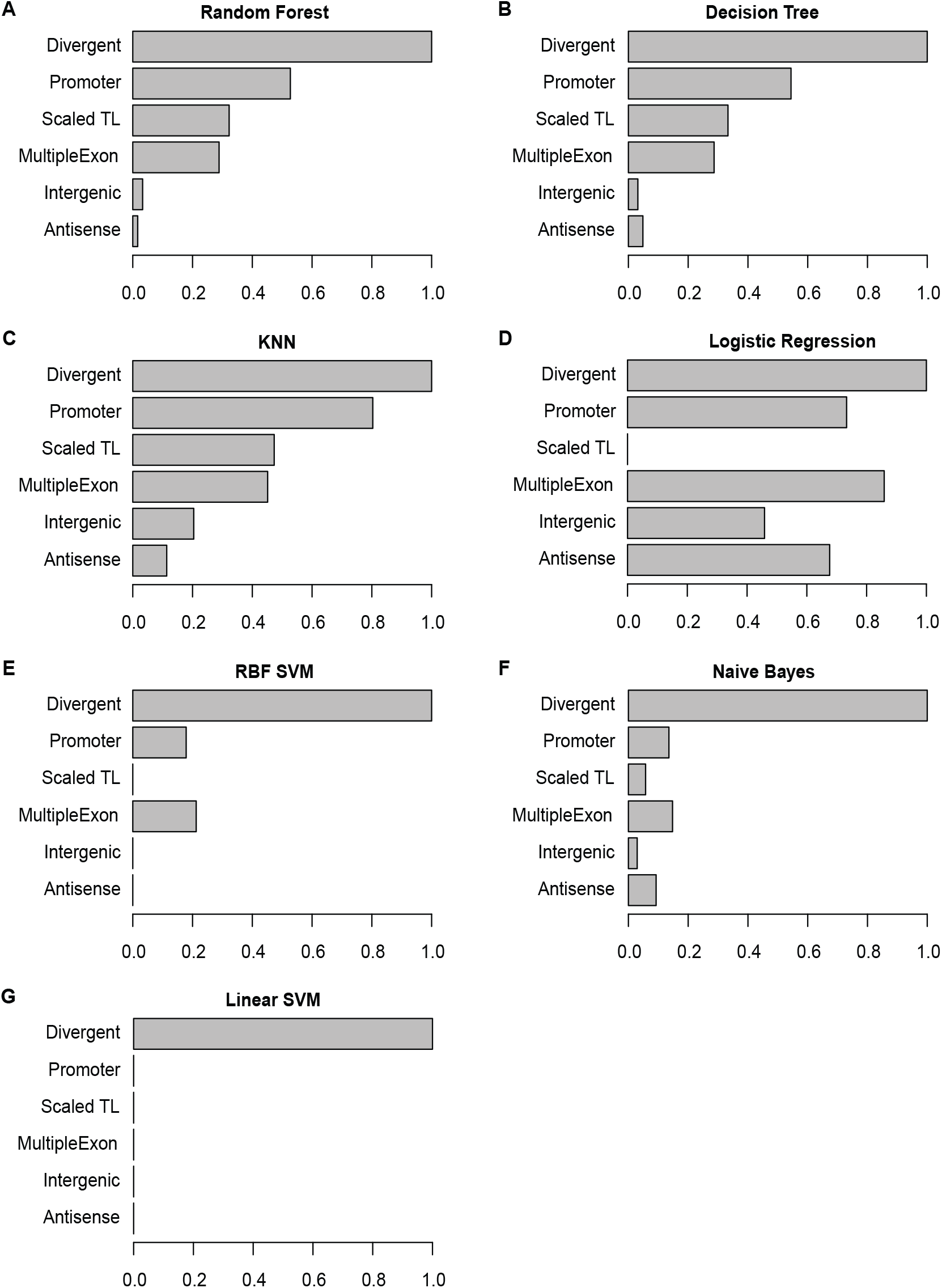
Feature importance for each of the seven final models. Each graph shows the scaled importance scores for each feature for the given model: random forest (A), decision tree (B), KNN (C), logistic regression (D), support vector machine (SVM) with RBF kernel (E), naïve bayes (F), and SVM model with linear kernel (G) models. For each model, the importance scores of all features were scaled to (0,1) by dividing the original score of the most significant feature (see Materials and Methods).

### *Flnc* achieves similar performance at the lncRNA gene locus level as at the transcript level

More than half (57%) of the true lncRNAs in the benchmark dataset have multiple exons. Among them, 43% of multi-exon lncRNA genes have undergone alternative splicing events, generating multiple lncRNA isoforms from the same lncRNA gene locus. *Flnc* has been trained and tested at the transcript level for each RNA-seq dataset. To examine the performance of *Flnc* at the locus level, we further tested *Flnc* on the lncRNA gene loci which encode only lncRNA transcripts. This test showed that *Flnc* can achieve similar performance at the lncRNA gene locus level as at the lncRNA transcript level (**Suppl Fig 5**), although *Flnc* performs better at the transcript level than at the locus level.

## DISCUSSION

We have developed a comprehensive pipeline and software *Flnc* which uses machine-learning models to accurately identify true lncRNAs from RNA-seq data. *Flnc* does not require matched transcription initiation profiles which are usually marked by H3K4me3 histone modifications. We trained the machine learning models on four types of features: transcript length, promoter signature, multiple exons, and genomic location, which show a high degree of discrepancy between true lncRNAs and false lncRNAs in our benchmark datasets. We benchmarked *Flnc* on both the transcript level, using all 46 datasets, and the dataset level, using the 46 RNA-seq datasets individually. *Flnc* can achieve up to 85% prediction accuracy using only RNA-seq data. This software improves of prediction of true lncRNAs, allowing their accurate prediction from samples, such as clinical samples, where ChIP-seq data is unavailable. *Flnc* can save time and resources by making the generation of H3K4me3 unnecessary and allowing the identification of lncRNAs from publicly available RNA-seq data.

The predictive power of *Flnc* relies on the quality of the benchmark datasets and the identification of true and false lncRNA within them; therefore, the prediction power of *Flnc* may be limited by factors that affect the reliability of the benchmark datasets. These factors include RNA-seq and ChIP-seq data quality, accuracy of transcript assembly and ChIP-seq peak calling, and sequencing depth. To minimize the effects of these factors, we examined the data quality of both RNA-seq and ChIP-seq data in the benchmark datasets. To improve the accuracy of transcript assembly, we integrated multiple transcript-assembly methods. It has been shown that the accuracy of ChIP-seq peak calling can be improved by using different approaches for long (≥70bp) and short (<70bp) reads studies (35, 51); therefore, we used this approach.

Compared to the true lncRNAs identified by H3K4me3 profiles, true lncRNAs predicted by *Flnc* (with random forest) are more likely to be divergent transcripts of nearby protein-coding genes and to have promoter signatures (**Suppl Fig 4B**). This result suggests that *Flnc* might overestimate divergent putative lncRNAs and putative lncRNAs with promoter signatures as true lncRNAs but underestimate the true lncRNAs that are in intergenic regions. One possible explanation for this is that divergent transcription and promoter signature are the two most important features for the three best models (random forest, decision tree and KNN).

Because different methods of generating and sequencing RNA-seq libraries result in slightly different collections of transcripts, feature importance may determine model performance for different types of RNA-seq data. We found that the random forest, decision tree and KNN models outperform than other models for stranded RNA-seq data generated from polyA-selected RNA (**Fig 3 and Suppl Fig 3**). In contrast, the linear SVM and naïve bayes models, which performed poorly for stranded polyA-selected RNA-seq data, achieve better prediction accuracy and precision than other models for rRNA-depleted and unstranded RNA-seq data. This may be because the random forest, decision tree and KNN models rely on three major features: divergent transcript, promoter signature, and transcript length (**Fig 6**), but the transcript length feature may not be applicable to nascent lncRNAs which are included in rRNA-depleted RNA-seq data. On the other hand, for unstranded RNA-seq data, the promoter signature feature may not applicable, as without accurate strand information, the promoters of these putative lncRNA cannot be inferred. Unlike the random forest, decision tree and KNN models, the linear SVM and naïve bayes models mainly rely on the divergent genomic structure feature (**Fig 6**), which is applicable for both nascent and unstranded lncRNAs.

*Flnc* includes seven embedded machine learning models (random forest, decision tree, logistic regression, and naïve Bayes, KNN, linear-SVM and RBF-SVM) which makes *Flnc* suitable for identifying lncRNAs from different types of RNA-seq data. Model performance depends somewhat on the type of RNA-seq data that is used as input. The best three models (random forest, decision tree and KNN) for the stranded polyA-selected RNA-seq data achieve very high consistency (93%-96%) in true lncRNA prediction (**Suppl Fig 3D**). Therefore, we recommend that users select any of these three models when using *Flnc* to identify lncRNAs from stranded polyA-selected RNA-seq data. Precision can be further improved by the ensemble approach (**Suppl Fig 3E**); therefore we recommend using the *ensemble* setting of the *Flnc* software, which provides users with the common set of true lncRNAs predicted by all models.

Although *Flnc* was trained on the benchmark datasets of lncRNAs identified from stranded polyA-selected RNA-seq data and matched H3K4me3 ChIP-seq data, the *Flnc* pipeline can be applied to identify lncRNAs from other types of RNA-seq data, such as stranded rRNA-depleted RNA-seq data and unstranded polyA-selected data. Based on the performance of the seven models on stranded rRNA-depleted RNA-seq data and unstranded polyA-selected data, we recommend using the linear KNN or naïve bayes for these two types of RNA-seq data.

The *Flnc* software can identify novel lncRNAs directly from RNA-seq data or evaluate if a transcript is a lncRNA or not. *Flnc* can take three types of input files, including raw RNA-seq data in the FASTQ format, transcript coordinates in the BED format, and transcript sequences in the FASTA format. For maximum portability and useability, *Flnc* is implemented in in Python 2 and the entire pipeline is wrapped within a Singularity container.

## DATA AVAILABILITY

The datasets analyzed in this study were downloaded from the NCBI GEO database: *https://www.ncbi.nlm.nih.gov/geo/*. The constructed benchmark dataset of true and false lncRNAs are available at *https://zhoulab.umassmed.edu/Flnc_data/*. The analyses were performed with Flnc version 1.0. The Flnc software is freely available on GitHub: *https://github.com/CZhouLab/Flnc*, along with a tutorial.

## SUPPLEMENTARY DATA

## ACKNOWLEDGEMENT

We thank Qian Qi for the helpful discussions about the ChIP-seq analysis. We thank Alan Mullen for review of the manuscript and insightful suggestions. We thank Feifan Liu for helpful discussion about the machine learning models. We thank Edith Pfister for revising the manuscript.

## SUPPLEMENTAL DATA FOR THE PAPER

## SUPPLEMENTAL METHODS

### Identification of putative lncRNAs

We upgraded the previous computational pipeline (**Fig 1A**) (Sigova et al., 2013; Zhou et al., 2016) with the latest tools to identify putative lncRNAs from raw RNA-seq data. The putative lncRNAs are transcripts longer than 200nt that lack coding ability. To identify the putative lncRNAs, we first assembled transcripts from raw RNA-seq data, then filtered out the transcripts that fall into any of the following categories: (i) transcripts with potential coding ability or other small noncoding RNAs, (ii) transcripts shorten than 200 nucleotide or with extremely low expression levels. Please see the details as follows.

1) ***Ab initio* assembly of transcripts from RNA-seq data**

We mapped each replicate of stranded polyA-selected RNA-seq data to the human reference genome (hg38/GRCh38) using HISAT2 v2.0.5 (Kim et al., 2015), and then assembled transcripts using StringTie v1.3.4 (Pertea et al., 2015) and Strawberry v1.1.2 (Liu and Dickerson, 2017).

The HISAT2 settings for single-end RNA-seq data were as follows:

*hisat2 -p 10 --dta -x < index of reference genome> -U < Reads*.*fastq > --add-chrname --rna-strandness <strandness> --fr --known-splicesite-infile <known splice site> --novel-splicesite-outfile <novel splice site file> --novel-splicesite-infile <novel splice site file> --seed 168 --phred33 --min-intronlen 20 --max-intronlen 500000*

The HISAT2 settings for paired-end RNA-seq data were as follows:

*hisat2 -p 10 --dta -x <index of reference genome> -1 <Reads_end1*.*fastq> -2 <Reads_end2*.*fastq>*

*--add-chrname --rna-strandness <strandness> --fr --known-splicesite-infile <known splice site> -*

*-novel-splicesite-outfile <novel splice site file> --novel-splicesite-infile <novel splice site file> -- seed 168 --phred33 --min-intronlen 20 --max-intronlen 500000*

The reference genes in GTF file format (genes.gtf) were downloaded from the Release 29 version of the GENCODE database (Harrow et al., 2012). We then assembled transcripts with the following settings of StringTie and Strawberry using HISAT2’s output bam file as input:

*stringtie <strandness> -p 20 -G genes*.*gtf -o d <output*.*gtf> -l ${1} -f 0 -m 200 -a 10 -j 1 -M 1 -g 50*

*<HISAT2_output_bam_file>*

*strawberry <strandness> -g genes*.*gtf -o <output*.*gtf> -p 20 -m 0 -t 200 -s 10 -d 50 --no-quant -- min-depth-4-transcript 0*.*1 <HISAT2_output_bam_file>*

Next, all transcripts assembled by either StringTie or Strawberry from all replicates were merged into one list through the StringTie merge function.

2) **Remove transcripts with potential coding abilities or other small noncoding RNAs**. First, we removed transcripts that overlapped with annotated protein-coding genes, pseudogenes, rRNAs, tRNAs, small nucleolar RNAs (snoRNAs), and microRNAs on the same strand. Second, we removed transcripts with protein coding potential. The coding potential of each remaining transcript was estimated by CPAT v1.2.4 (Wang et al., 2013), LGC v1.0 (Wang et al., 2019), PLEK v1.2 (Li et al., 2014), and CPPred (Tong and Liu, 2019). We used the default threshold or the suggested threshold of each tool to determine the coding abilities for each of the remaining transcripts. Third, we removed any remaining transcripts that overlapped on the same strand with the transcripts removed in the previous two steps.

3) **Remove short and lowly expressed transcripts**

We removed the remaining transcripts that were shorter than 200nt. Then, we removed the remaining transcripts that are lowly expressed. A transcript is considered lowly expressed if it had either 1) less than ten reads per transcript, or 2) its expressed level is less than the smallest FPKM peak value that occurs in the bottom half of the FPKM distribution curve for all remaining expressed transcripts.

## SUPPLEMENTAL FIGURES

**Supplementary Figure 1:**
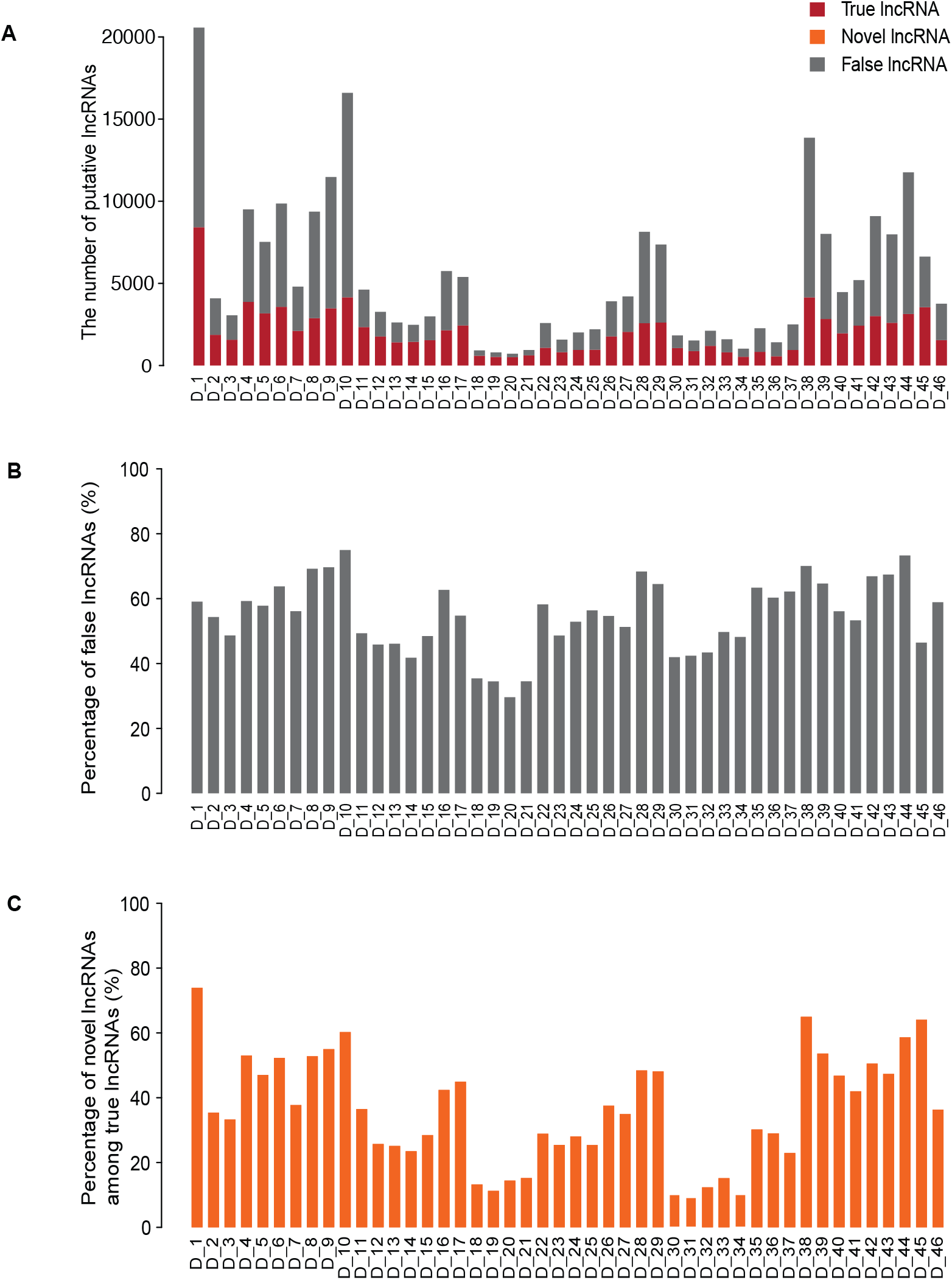
True and false lncRNAs in each of the 46 benchmark datasets. The number of putative true (red) and false (grey) lncRNAs (A) and the percentage of false lncRNAs in each dataset (B). The putative true lncRNAs include a high percentage of novel lncRNAs in each dataset (C).

**Supplementary Figure 2:**
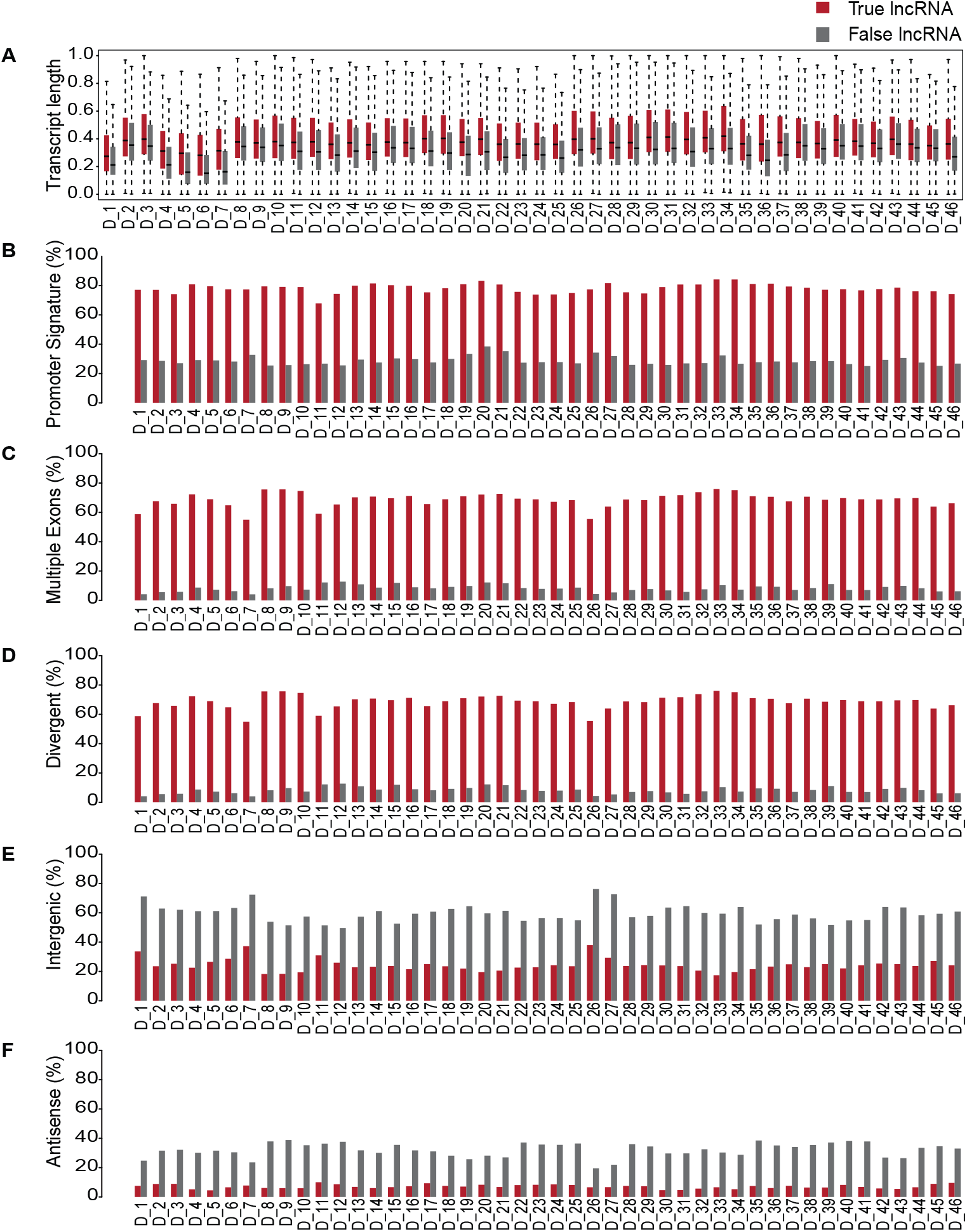
Genomic features of true and false lncRNAs in each of the 46 benchmark datasets. In each of the 46 benchmark datasets, true and false lncRNAs can be distinguished by (A) transcript length (A), the presence of an upstream promoter signature (B), the presence of multiple exons (C). True lncRNAs are more often divergently transcribed from the promoters of protein-coding genes (D), and less likely to be intergenic (E), or antisense to protein-coding genes (F). For (A), the boxplots represent the scaled transcript lengths of true lncRNAs and false lncRNAs across each of the 46 benchmark datasets. The error bars are the 95% confidence interval, the bottom and top of the box are the 25th and 75th percentiles, the line inside the box is the 50th percentile (median). For (B-F), each bar-plot represents the percentage of a feature among the true and false lncRNAs across each of the 46 benchmark datasets.

**Supplementary Figure 3:**
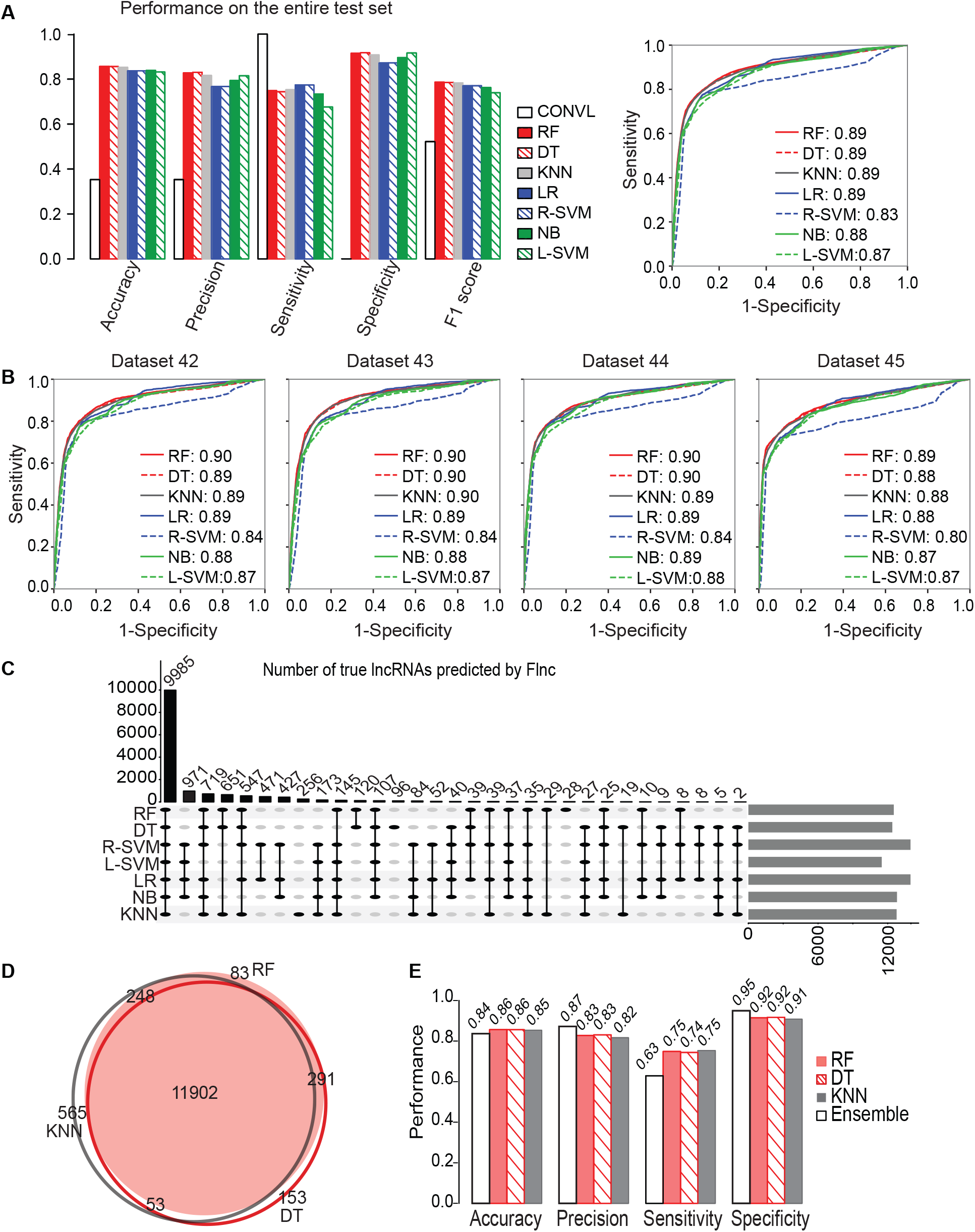
Performance of the seven final machine learning models used in *Flnc*. (A) The seven final models of *Flnc* outperformed the conventional method (without models) on the entire test set. *The left bar graph shows the performance metrics accuracy, precision, sensitivity, specificity and F1 score. The right graph shows the receiver operating characteristic (ROC) curve for each model. Area Under the ROC curve (AUROC) score is shown next to each ML model*. (B) The ROC curves on the dataset 42-45 within the test set. (C) The upset plot shows the number of true lncRNAs commonly predicted by different models. The vertical bars in black represent the number of true lncRNAs commonly predicted by the models that are highlighted and connected by black line below the bar. For example, 9985 true lncRNAs are commonly predicted by all the seven models in the test set; and 971 true lncRNAs are commonly predicted by RBF SVM, linear SVM, logistic regression and naïve Bayes models. The horizontal bars in gray represent the number of true lncRNAs predicted by each model listed on the left side. (D) The Venn diagram of the predicted true lncRNAs by the three best models (random forest, decision tree and KNN). (E) Comparison of the performance of the three best models and the ensemble approach (in white). The result predicted by ensemble approach will improve the prediction precision and specificity with the cost of a reduced sensitivity. *Abbreviations: CONVL, conventional approach; RF, random forest; DT, decision tree; R-SVM, RBF support vector machine; L-SVM, linear support vector machine; LR: linear regression; NB, naïve bayes; KNN*, k-nearest neighbors.

**Supplementary Figure 4:**
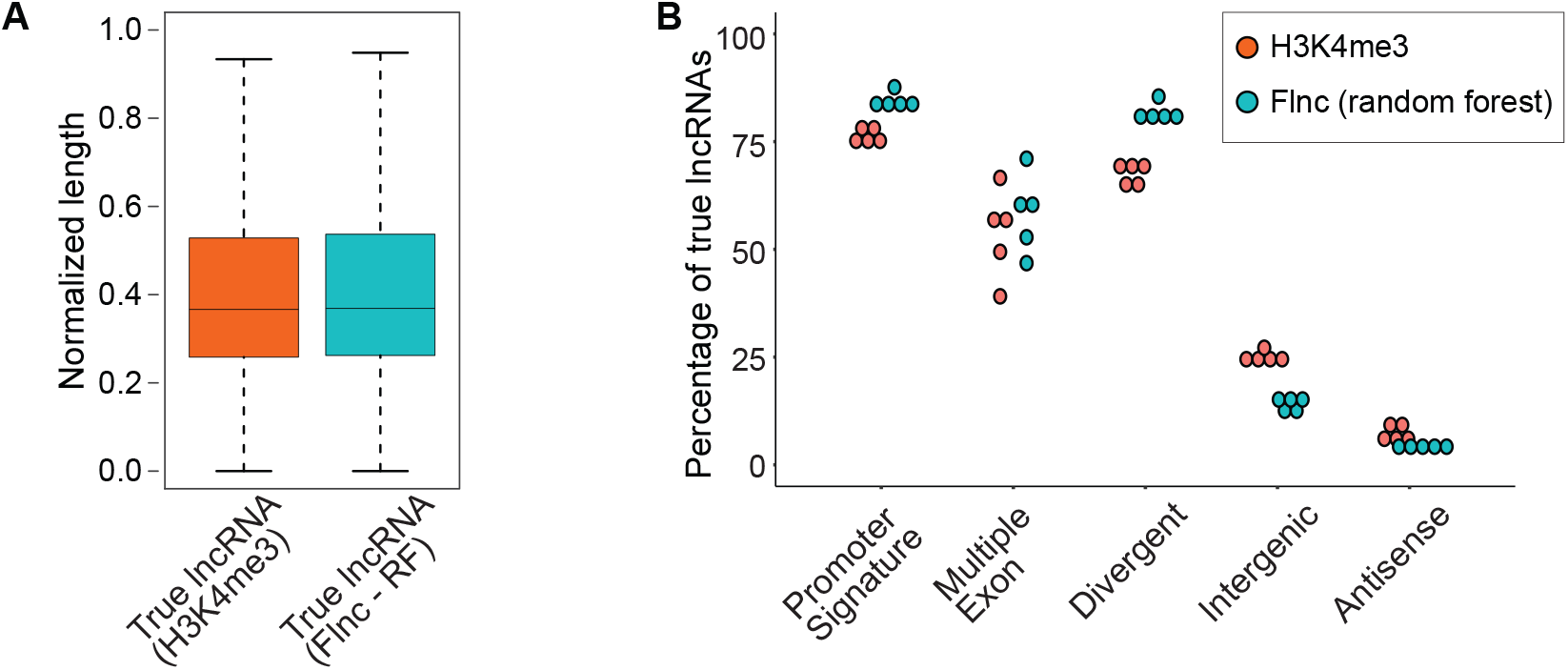
The genomic features of true lncRNAs predicted by *Flnc* and true lncRNAs determined by H3K4me3 profiles in the test set. (A) The true lncRNAs predicted by *Flnc* (with random forest model) show similar normalized transcript length distribution as the true lncRNAs determined by H3K4me3 ChIP-seq data. (B) The true lncRNAs predicted by *Flnc* (with random forest model) include significantly more divergent transcripts and transcripts with promoter signatures than the true lncRNAs determined by H3K4me3 ChIP-seq data, whereas multiple exon features exhibit similar percentage between these two groups of lncRNAs.

**Supplementary Figure 5:**
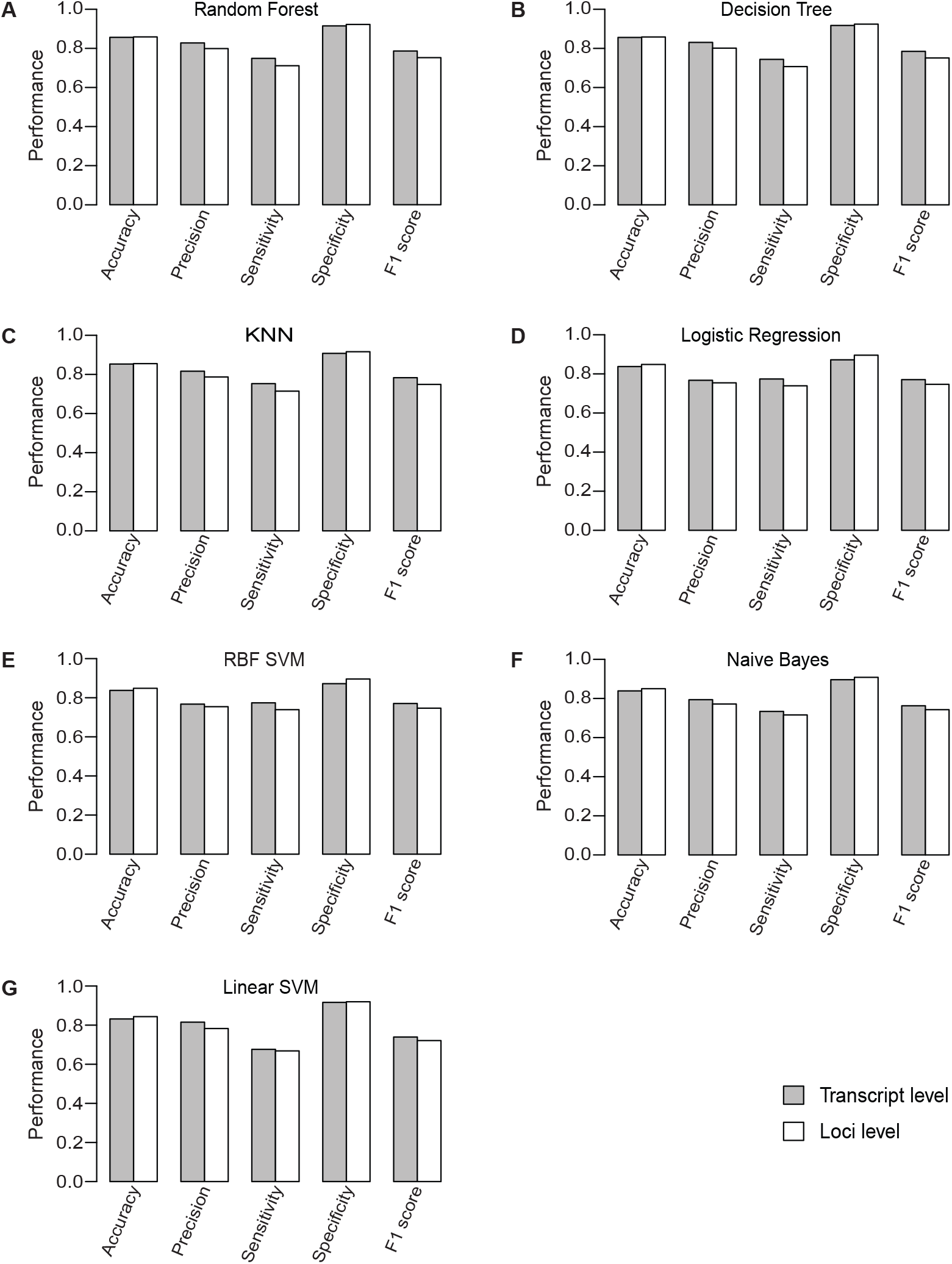
*Flnc* achieves similar performance at the lncRNA gene locus level as at the transcript level. Each graph shows the performance for the given model: random forest (A), decision tree (B), KNN (C), logistic regression (D), support vector machine (SVM) with RBF kernel (E), naïve bayes (F), and SVM model with linear kernel (G) models.

